# Post-catalytic spliceosome structure reveals mechanism of 3'-splice site selection

**DOI:** 10.1101/212811

**Authors:** Max E. Wilkinson, Sebastian M. Fica, Wojciech P. Galej, Christine M. Norman, Andrew J. Newman, Kiyoshi Nagai

## Abstract

Introns are removed from eukaryotic mRNA precursors by the spliceosome in two transesterification reactions – branching and exon ligation. Following branching, the 5'-exon remains paired to U5 snRNA loop 1, but the mechanism of 3'-splice site recognition during exon ligation has remained unclear. Here we present the 3.7Å cryo-EM structure of the yeast P complex spliceosome immediately after exon ligation. The 3'-splice site AG dinucleotide is recognised through non-Watson-Crick pairing with the 5'-splice site and the branch point adenosine. A conserved loop of Prp18 together with the α-finger and the RNaseH domain of Prp8 clamp the docked 3'-splice site and 3'-exon. The step 2 factors Prp18 and Slu7 and the C-terminal domain of Yju2 stabilise a conformation competent for 3'-splice site docking and exon ligation. The structure accounts for the strict conservation of the GU and AG dinucleotides of the introns and provides insight into the catalytic mechanism of exon ligation.

---

Pre-mRNA splicing is catalysed by a dynamic molecular machine called the spliceosome (*1*), which employs a single, RNA-based, active site (*2, 3*) to catalyse two sequential transesterification reactions that excise non-coding introns from pre-mRNA and ligate the coding exons to form mature mRNA. Introns are marked by GU and AG dinucleotides at the 5'- and 3'-splice sites (SS) respectively and a branch point (BP) adenosine upstream of the 3'SS: these nucleotides are invariant except in introns removed by the metazoa-specific minor spliceosome (*4*). The spliceosome assembles *de novo* on each pre-mRNA by the ordered joining of five small nuclear ribonucleoprotein particles (snRNPs) and the NineTeen and NineTeen-Related (NTC and NTR) protein complexes, along with additional protein factors (*1*). First the U1 and U2 snRNPs recognise the 5'SS and BP sequences of the pre-mRNA respectively, forming A complex. Next the pre-assembled U4/U6.U5 tri-snRNP joins to form pre-B complex, followed by spliceosome activation via B complex to form B^act^ complex. Subsequent Prp2-mediated remodelling yields B^*^ complex, which is competent to carry out the first transesterification reaction, called branching, when the 2'-hydroxyl of the conserved BP adenosine in the intron performs a nucleophilic attack on the 5'SS, producing the cleaved 5'-exon and a branched lariat-intron intermediate. The resulting C complex is then remodeled to C* complex upon dissociation of step 1 (branching) factors by the DEAH-box ATPase Prp16 (*5, 6*). In C* complex, step 2 (exon ligation) factors promote docking of the 3'SS into the active site (*7*) and exon ligation, via nucleophilic attack of the 3'-hydroxyl of the 5'-exon on the 3'SS. The resulting P complex contains the ligated exons (mRNA) and the excised lariat-intron. The newly formed mRNA is then released from P complex by the DEAH-box ATPase Prp22 (*8*), forming the intron-lariat spliceosome (ILS) which is disassembled by the DEAH-box ATPase Prp43 (*1*) to recycle the snRNPs for further rounds of splicing.

Recent cryo-electron microscopy (cryo-EM) studies of yeast (*6, 9-15*) and human spliceosomes (*16, 17*) have elucidated the configuration of the RNA-based active site and many mechanistic details of splice site recognition and catalysis, as well as the role of specific protein factors (*18*). Structures of C complex (*6, 12*) showed how the spliceosome recognises and positions the 5'SS and BP sequences in the active site through RNA-RNA interactions with U6 snRNA and U2 snRNA, while the branching factors Cwc25, Yju2, and Isy1 lock these sequences in a conformation competent for catalysis. Structures of C and C* complex (6, *14-17*) revealed how Prp16 remodels the spliceosome into the exon ligation conformation, which is stabilised by the exon ligation factors Prp18 and Slu7 (*19, 20*). In both C and C* complexes the 5'SS and 5'-exon remain paired with U6 snRNA (*21, 22*) and loop 1 of the U5 snRNA (*23, 24*), respectively. However in the C* complex structures the 3’-exon and 3’SS are not yet docked into the active site. Thus it has remained unclear how the spliceosome selects and docks the 3'SS while aligning the 3'-exon for step 2 catalysis. Additionally, it was not known how Slu7 and Prp18 interact with the 3'SS to promote docking.

Here, we report the cryo-EM structure of the *Saccharomyces cerevisiae* post-catalytic P complex at near-atomic resolution, showing the catalytic step 2 configuration of the active site. The structure reveals how the 3'SS and 3'-exon are recognised by the spliceosome and shows the critical role of the branched lariat-intron in promoting the chemistry of exon ligation.

## Overall architecture of P complex

We produced P complex by an *in vitro* splicing reaction supplemented with dominant-negative mutant Prp22 protein to prevent mRNA release from the spliceosome (*25*). Complexes with a docked 3'-exon were selectively purified via MS2-MBP fusion protein (see Methods). The resulting spliceosomes contained spliced mRNA and excised intron and were enriched in Prp22, as expected (**Fig. S1**). We obtained a cryo-EM reconstruction at 3.7Å overall resolution and modelled 45 components (**Fig. S2, Tables S1-2**)

P complex has the same overall exon ligation conformation as C* complex (**Fig. 1A**). Relative to the branching conformation of C complex (*6, 12*), the branch helix between the intron and U2 snRNA has rotated 75 degrees out of the active site, extracting the BP adenosine and creating space for the incoming 3'-exon in the active center (**Fig. 2A,B**). Branch helix rotation is accompanied by movement of the U2 snRNP and its attached NTC protein Syf1, and Prp16-mediated release of the branching factors Cwc25, Isy1 and the N-terminal domain of Yju2 (*14, 15*) (**Fig. S4A**). The undocked branch helix is stabilised in its new position by the WD40 domain of Prp17, and by the Prp8 RNaseH domain, which has rotated to insert its β-hairpin near the BP (*14, 15*). The exon ligation factors Prp18 and Slu7 occupy the same locations as they do in C* complex, with the α-helical domain (*26*) of Prp18 binding the outer surface of the Prp8 RNaseH domain (**Fig. 1C**). While in C* complex we were only able to assign the C-terminal globular domain of Slu7, the higher quality density in P complex allowed us to essentially complete this model, revealing a sprawling architecture that spans 120Å of the spliceosome (**Fig. 4A**). Prp22, which replaces Prp16 in both C* and P complex, is stabilised onto the Prp8 N-terminal domain through interactions with the C-terminus of Yju2.

**Figure 1.**
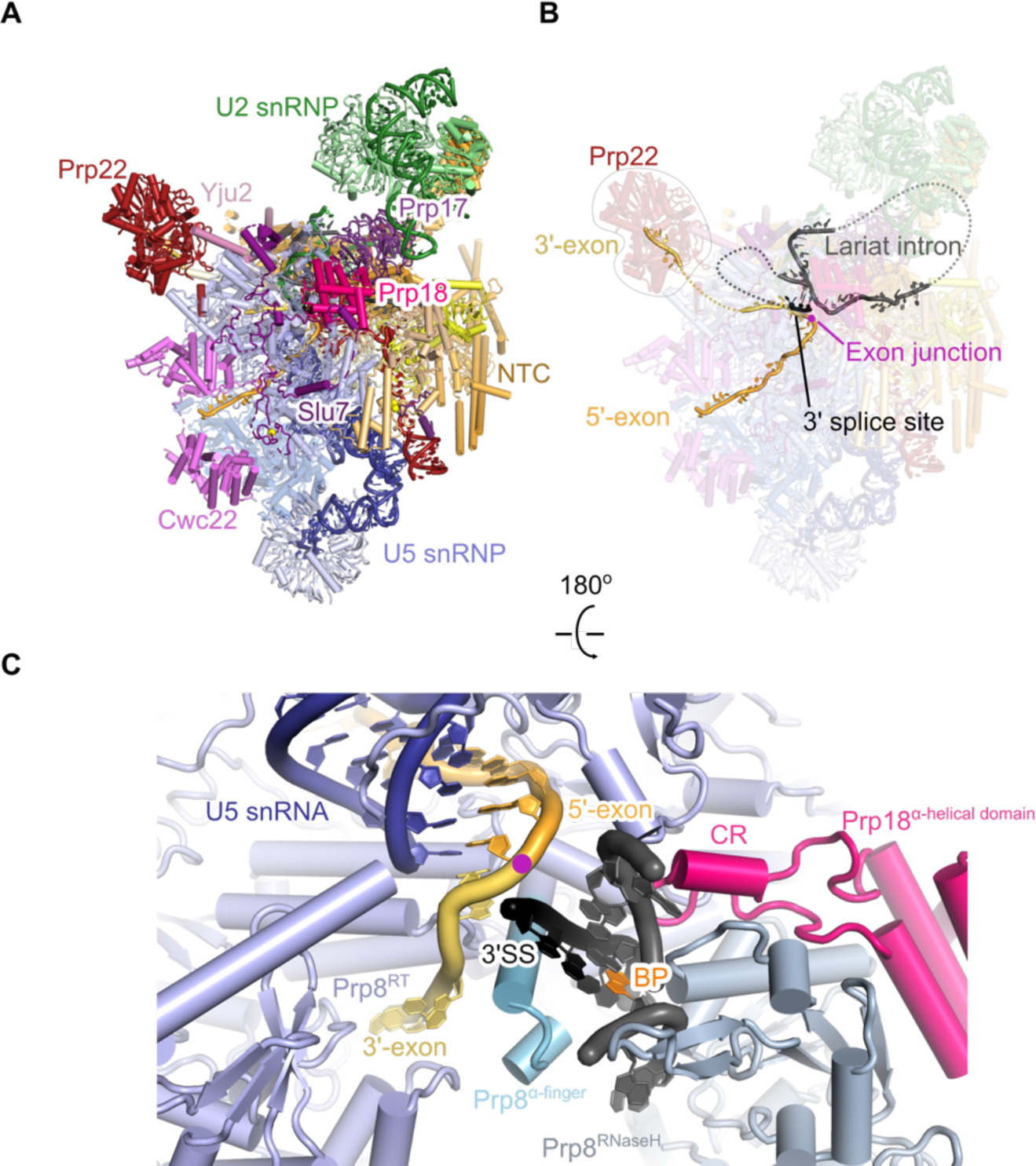
P complex structure. **A**, Overview of the P complex spliceosome. NTC, NineTeen Complex. **B**, The same view of P complex with the path of the substrate intron and exons shown. Dotted lines indicate the path of nucleotides not visible in the density. **C**, Binding of substrate at the core of P complex. U2 and U6 snRNAs and NTC/NTR proteins are omitted for clarity. 3'SS, 3' splice site; RT, Prp8 reverse transcriptase domain; CR, Prp18 conserved region.

## RNA active site

As observed in C* complex, the RNA catalytic core of the spliceosome is similar between branching and exon ligation conformations (**Fig. 2A,B**). The catalytic triplex formed by U2 and U6 snRNAs, harboring the catalytic metal ions, is unaltered (*2, 3, 27*), and the 5'-exon remains base-paired to loop 1 of U5 snRNA snRNA (*23, 24*). The RNA catalytic core of the P complex spliceosome remains essentially unchanged compared to C* complex (*14-17*), except that the 3'-exon and the intron 3'SS are now accommodated in the active site (**Figs. 2B, 1B,C**). Our P complex map clearly shows that the 3'-exon is ligated to the 5'-exon and that the 3'SS is docked into the space occupied by the branch helix in C complex (**Figs. 2B, S3**). The position of the 3'SS relative to the 3'-exon suggests that prior to exon ligation, the pre-mRNA undergoes an almost 180° turn to expose the 3'SS scissile phosphate. This deformation is similar to that seen during branching, when the 5'SS is highly bent to expose the scissile phosphate to the branch-point adenosine nucleophile (*6, 12*) (**Fig. S5**).

**Figure 2.**
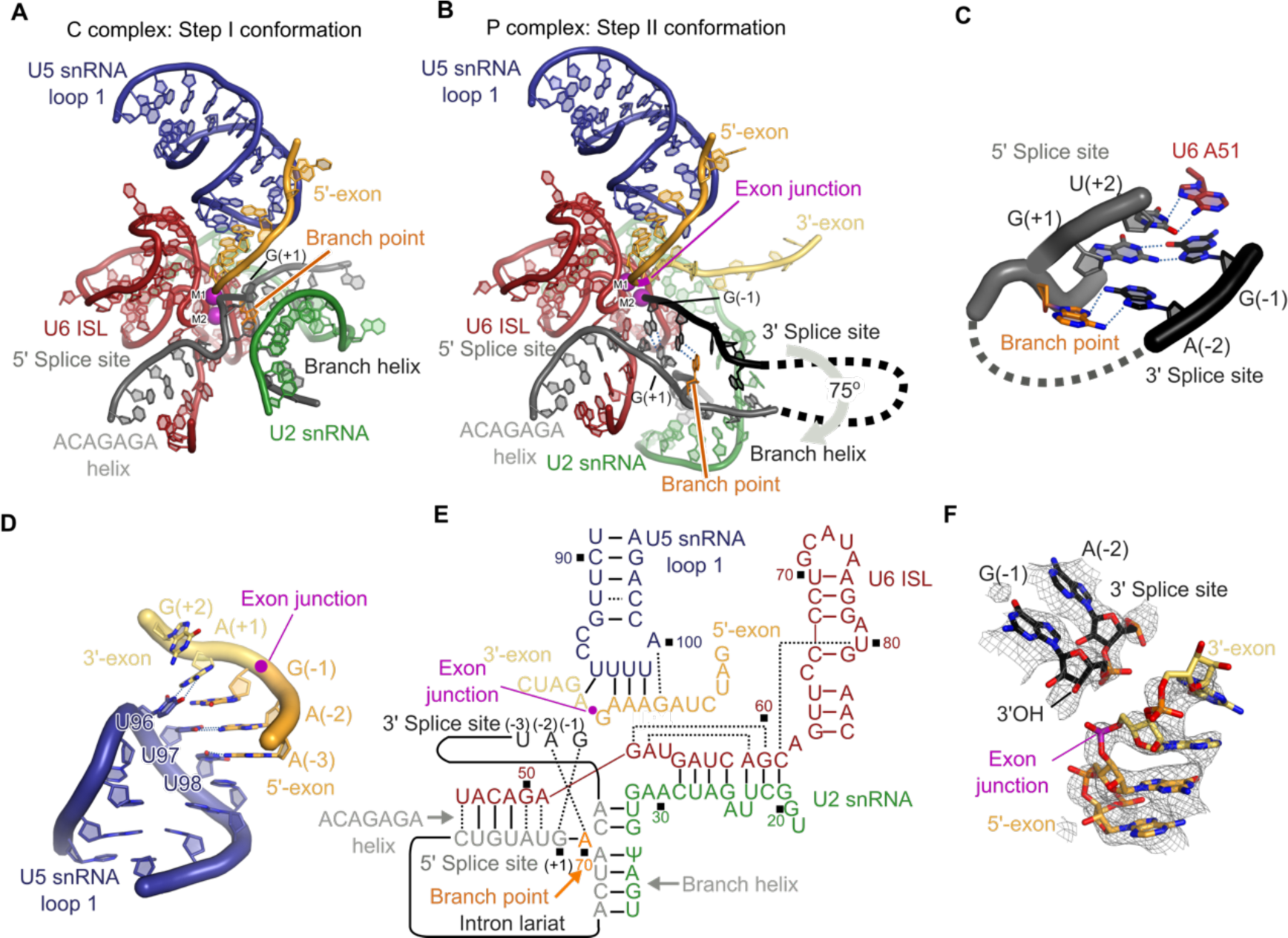
Structure of the RNA catalytic core. **A**, Key RNA elements at the active site of the C complex spliceosome. ISL, internal stem-loop; M1 and M2, catalytic metal ions one and two. **B**, Equivalent view to **A** of the active site of the P complex spliceosome. M1 was not visible in the density and its position is inferred from C complex. **C**, Non-Watson-Crick RNA-RNA interactions mediate recognition of the 3' splice site. Putative hydrogen bonds are shown with dotted blue lines. BP, branch point adenosine. **D**, U5 snRNA loop 1 aligns the 5'- and 3'-exons for ligation. U5 snRNA U96 is flexible and can pair to both G(-1) of the 5'-exon and A(+1) of the 3'-exon. **E**, Base-pairing scheme of the P complex active site. Watson-Crick pairing is indicated with lines, other base pairs with dotted lines. Ψ, pseudo-uridine. **F**, CryoEM density around the exon junction for the 5'-exon, 3'-exon and 3' splice site.

The first two bases of the 3'-exon are well ordered and extend the 5'-exon with A-form helical geometry and regular base stacking (**Fig. 2D**) whereas ten nucleotides of the 5'-exon are well ordered in the channel between the N-terminal and Large domain of Prp8. Density for the 3'-exon downstream of G(+2) becomes weaker and follows the surface of the Prp8 reverse transcriptase (RT) domain up towards Prp22 (**Fig. S3F**), consistent with crosslinking experiments (*25*). This arrangement is consistent with the role of Prp22 in pulling the ligated exon from the 3' direction to release mRNA. Our C complex structure revealed a similar mechanism of remodeling by Prp16 in pulling the intron (6, 18). It was previously shown that U5 snRNA loop 1 aligns the 5'-exon and 3'-exon for ligation (*23, 24*). Indeed, U5 snRNA U96, which points away from the active site in B^act^, C, and C* complexes, can pair with A(+1) of the 3'-exon in P complex (**Fig. 2D**). Thus the ends of both the 5'-exon (nucleotides −2 to −4) and the 3'-exon (nucleotide +1) base-pair with U5 snRNA loop 1, explaining previous genetic and crosslinking data (*23, 24*).

In yeast the intron sequence of the 3'SS immediately preceding the 3'-exon is stringently conserved as Y(-3)A(-2)G(-1), where Y is any pyrimidine (*28*) (**Fig. S6A**). In our P complex structure the phosphodiester bond at the 3'SS is cleaved, but the 3'-hydroxyl of the 3'SS nucleotide G(-1) remains close to the phosphate of the newly-formed exon-junction, consistent with observations that exon ligation is reversible (*29*) (**Fig. 2F**). This suggests that our structure represents the state of the spliceosome immediately after exon ligation, allowing us to infer the mechanism of 3'SS recognition (**Fig. 2C**). Remarkably, the Hoogsteen edge of the 3'SS G(-1) forms a base pair with the Watson-Crick edge of the 5'SS G(+1), while stacking on U6 snRNA A51, which remains paired to U(+2) of the 5'SS as in C* complex (*14*). This arrangement allows the 3'-hydroxyl of 3'SS G(-1) to project towards the active site (**Fig. 2F**). The Hoogsteen edge of the 3'SS A(-2) interacts with the Hoogsteen edge of the branch point adenosine, which is still linked via its 2'-hydroxyl to the 5'SS G(+1). Thus, 3'SS recognition is achieved through RNA base-pairing with the 5'SS and the BP adenosine. This mechanism is consistent with the genetic interactions between the first and last bases of the intron (*30, 31*) and exon ligation defects observed in branch point mutants (*32, 33*) (**Fig. S6**). We additionally observed ordered density for 3'SS bases (-3) to (-5), while the 20 bases connecting the 3'SS to the branch helix were not visible and are likely disordered as they would loop out of the spliceosome from the branch helix and back into the spliceosome just upstream of the 3'SS.

In our cryo-EM map we observed density consistent with the presence of the catalytic Mg^2+^ ion M2 identified by metal rescue studies, which is chelated by phosphate oxygen ligands from U6 snRNA bases U80, A59 and G60 (*2*). Density for M1 is not visible, consistent with P complex being in a post-catalytic state, whereas M1 was predicted to coordinate the nucleophile in the pre-catalytic state and density consistent with M1 was indeed visible in C* complex (*14*) (**Figs. S3,S5**).

## Proteins around the active site

Compared to C* complex, which lacked the docked 3'-exon and 3'SS, new protein density becomes visible around the P complex active site. Prp8 and Prp18 cooperate to stabilise the docked 3'-splice site as well as the binding of the 3'-exon. The pairing between A(+1) of the 3'-exon and U96 of U5 snRNA loop 1 is sandwiched between the α-finger and the reverse transcriptase domains of Prp8 (**Fig. 1C**) (*34*). The α-finger (residues 1565-1610) of Prp8 forms an extended helix that wedges between the 3'-exon and 3'SS (**Fig. 3B, C, Fig S4**). Before exon ligation, the connected 3'-exon and 3'SS would wrap around this helix, exposing the 3'SS phosphate for attack by the 5'-exon (**Fig. 3B, Fig. S5**). Alpha-finger residue Arg1604, which is essential for exon ligation (**Figs. 3D, S8**), contacts the phosphate backbone of the 3'SS residues −3 and −4. The highly conserved Gln1594 forms a hydrogen bond with the O2 carbonyl group of U(-3), and cytosine at the (-3) position could form the same hydrogen bond explaining the preference for pyrimidines at (-3) of the 3'SS (**Figs. 3C, S6**). The non-Watson-Crick RNA pairing between the BP, the 5'SS, and the 3'SS is reinforced by these sequences being clamped between the Prp8 α-finger and the β-hairpin of the Prp8 RNaseH domain (**Fig. 3A**). Consistent with the structure, mutations in the β-hairpin suppress the exon ligation defects of mutations at 3'SS A(-2) and the BP (*35*), further underscoring the essential structural role of Prp8 in 3'SS docking. The so-called conserved region of Prp18 (*26*) forms a loop that penetrates from outside the spliceosome through a channel formed by Prp8 into the active site, where it buttresses against nucleotides U(-3) and C(-4) of the 3'SS (**Figs. 1C, 3A,B**). Deletion of this conserved loop in Prp18 affects 3'SS selection (*36*), consistent with its direct involvement in exon ligation.

**Figure 3.**
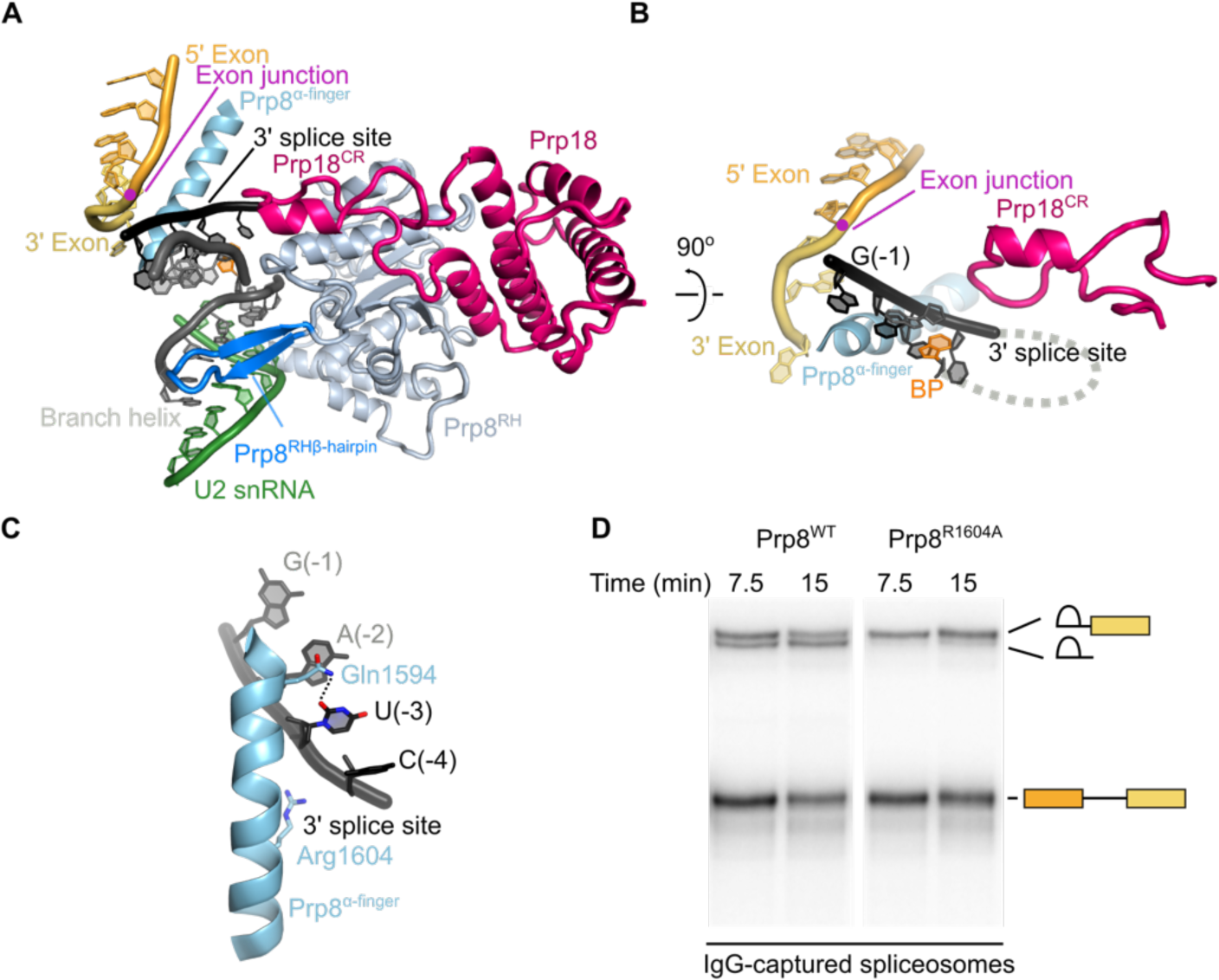
Proteins at the active site. **A**, The Prp8 α-finger and β-hairpin clamp around the active site, with Prp18 bound on the outer face of the Prp8 RNaseH domain. CR, Prp18 conserved region; RH, Prp8 RNaseH domain. **B**, The Prp8 α-finger contacts both 3'-exon and 3' splice site; Prp18 CR loop contacts the 3'SS from the opposite side. **C**, Residues of the Prp8 α-finger that contact the 3' splice site. **D**, In vitro splicing reaction with wild-type (WT) and Arg1604 to Ala (R1604A) mutant Prp8. RNA species found in Prp8-immunoprecipitated spliceosomes are labelled schematically. R1604A causes a second-step defect, evidenced by accumulation of lariat intron-3'-exon intermediate.

## Step 2 factors promote exon ligation

In C complex the branching factors Cwc25, Isy1, and Yju2 make extensive contacts with the branch and ACAGAGA helices and together directly stabilise the docking of the distorted branch helix into the active site to allow efficient branching (*6, 12*). In P complex, the exon ligation factors Prp18 and Slu7 make only one direct contact with the active site RNA, via the Prp18 conserved loop (**Fig. 3A, B**), while the α-helical domain of Prp18 binds the outer face of the Prp8 RNaseH domain (**Fig. 4A**). The N-terminus of Slu7 binds to the Cwc22 C-terminal and Prp8 linker and Prp8 endonuclease domains. Slu7 then descends to the Prp8 N-terminal domain, where it is anchored by its zinc-knuckle domain before ascending and passing through the interface between the Prp8 N-terminal and endonuclease domains. After a helical interaction with Prp18 the globular C-terminal region of Slu7 binds the inner surface of the Prp8 RNaseH domain (**Fig. 4A**). This mostly peripheral binding of exon ligation factors suggests they promote splicing by a less direct mechanism than the branching factors which lock the branch helix in the active site.

**Figure 4.**
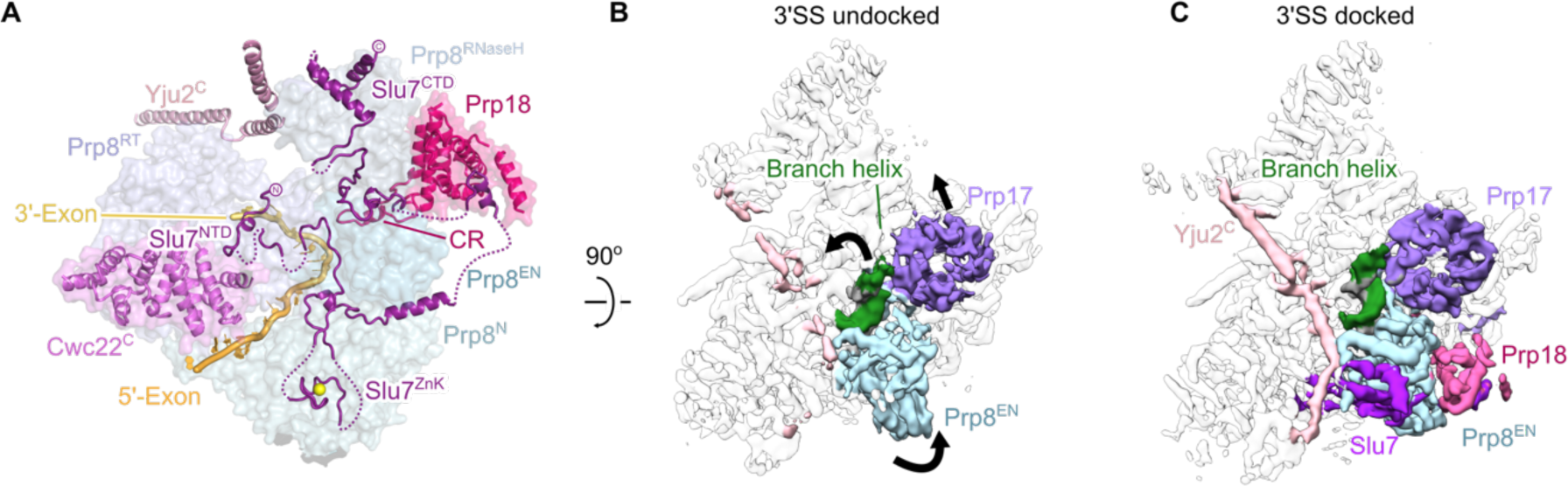
Docking of the 3' splice site is associated with binding of exon ligation factors. **A**, The binding of exon ligation factors Slu7 and Prp18 to the surface of P complex. Disordered segments of Slu7 are shown with dotted lines. Domains of Prp8 are coloured as indicated. N, Prp8 N-terminal domain; RT, Prp8 reverse transcriptase domain; EN, Prp8 endonuclease domain; ZnK, Slu7 zinc knuckle domain; CR, Prp18 conserved region. **B, C**, Cryo-EM density maps for P complex with the 3'SS docked and undocked. Maps were filtered to 5 Å resolution to aid visualization. Movements of the branch helix, Prp17, and the Prp8 endonuclease domain are indicated.

## Docked and undocked conformations of P complex

We performed global classification of our cryo-EM dataset to assess the conformational dynamics of P complex spliceosomes. A subset comprising approximately half of the purified P complex particles entirely lacks density for the 3'SS and Prp8 α-finger (**Fig. S7B**). These particles in the “undocked” conformation also lack density for Prp18 and Slu7, indicating that the presence of exon ligation factors correlates with stable docking of the 3'SS (**Fig. 4B,C**). This is consistent with previous biochemistry and genetics which suggest that Slu7 and Prp18 act after Prp16-mediated remodelling (*7, 37*) and promote juxtaposition of the splice sites in the exon ligation conformation (*7*). In the “undocked” conformation the junction between the 5'-exon and 3'-exon in ligated mRNA is still visible, confirming the undocked conformation represents a P complex state. The branch helix, which is locked in place by exon ligation factors in the docked conformation, undergoes slight movement together with Prp17 in the undocked conformation, suggesting that it is more flexible in the absence of exon ligation factors (**Figs. 4B, C, S7**). Because the BP adenosine in the branch helix and the attached 5'SS G(+1) form the recognition platform for the 3'SS AG sequence, it is likely that the 3'SS can only stably dock when the branch helix is held rigid in the docked conformation.

Interestingly, the docked conformation competent for exon ligation is also associated with stronger density for two long collinear alpha helices spanning the width of the spliceosome from the C-terminus of Syf1 to the Prp8 RNaseH domain and contacting Cef1 (**Fig. 4C**). This density was previously seen weakly in C* complex (*14*) but limited local resolution precluded its assignment. The higher quality of the P complex density allowed assignment of these helices as the C-terminus of Yju2. In contrast, the N-terminus of Yju2 binds onto the branch helix and acts as a branching factor in C complex. However, the Yju2 N-terminus is no longer visible in C* or P complexes. Previous experiments showed that the N-terminal and C-terminal domains of Yju2 can act in trans; the N-terminal domain is essential for branching but impedes exon ligation, while the C-terminal domain promotes exon ligation, despite not being essential for viability (*38*). Our structure explains these apparently opposing roles of Yju2 and suggests the C-terminal domain stabilizes binding of Prp22 and Slu7 and acts as a brace, further restricting flexibility in the docked conformation. Thus Yju2 is both a branching and exon ligation factor and Prp16-dependent remodelling of C complex effects an exchange of its stable binding to the spliceosome from the N-terminal to the C-terminal domain.

The two alternative forms of P complex suggest that exon ligation factors aid exon ligation in part by stabilising the docked conformation: Prp18 and Slu7 bind together to the Prp8 RNaseH domain in its rotated conformation induced by Prp16 remodelling of C complex, while Slu7 anchors the RNaseH domain in place via multi-pronged interactions with the other domains of Prp8 (**Fig. 5**). Slu7 may also promote exon ligation by additional mechanisms. Slu7 is dispensable for exon ligation when the distance between the BP and 3'SS is less than nine nucleotides (*19, 39*), consistent with the seven ordered nucleotides we see in P complex, which could be joined by an additional two nucleotides. The intron region between the BP and the docked 3'SS would likely project outward from the spliceosome through an opening between the Prp8 RNaseH, Linker, and RT domains. Intriguingly, the N-terminus of Slu7 binds at the base of this opening (**Figs. 4A, 1B**) and truncation of the N-terminus abolishes the ability of Slu7 to promote exon ligation for pre-mRNAs with long BP-3'SS distances (*37*). Thus the N-terminus of Slu7 could influence the entropic cost of 3'SS docking. In conjunction with Prp18, Slu7 could also determine the direction from which the 3'SS docks into the active site and could thus promote the correct topology for stable 3'SS docking.

**Figure 5.**
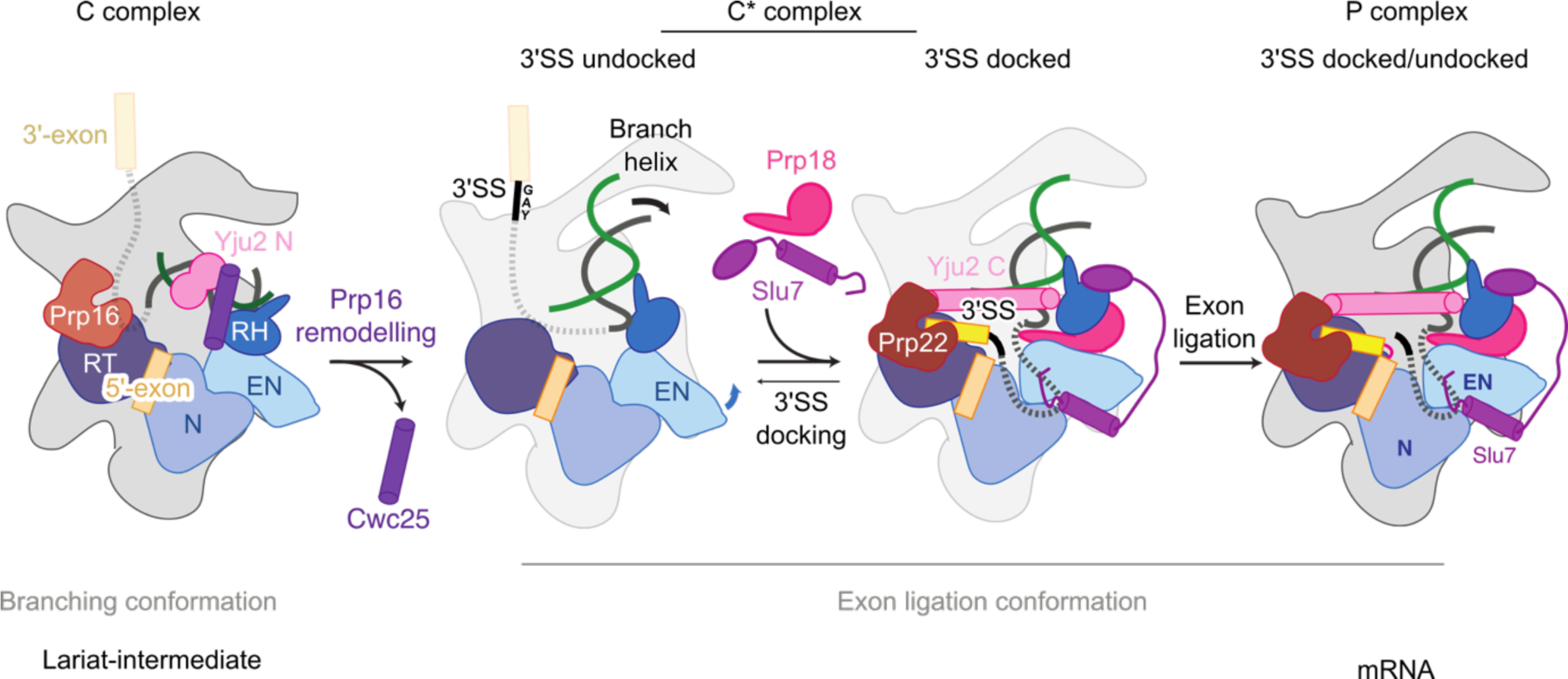
Model for the action of exon ligation factors. Following Prp16-mediated remodelling of C complex the branching factors Cwc25, Yju2 N-domain and Isy1 (not depicted) are removed. The undocked branch helix is then locked in a conformation competent for second step catalysis by the binding of exon ligation factors Prp18 and Slu7 and the C-domain of Yju2. The 3'SS docked and undocked conformations may be in equilibrium due to flexibility of the branch helix and Prp8 Endonuclease domain in the C* conformation.

## Conclusions

The molecular mechanism of 3'SS recognition during the catalytic phase of pre-mRNA splicing had been elusive despite decades of functional studies. Our P complex structure now shows that the 3'SS is recognised by pairing with the 5'SS and the branch point adenosine. This interaction involves all invariant nucleotides (GU and AG) at the 5' and 3' ends of the intron and the invariant branch point adenosine, making the exon ligation step a final quality check of the splicing reaction. Indeed, mutation of any of these invariant nucleotides impairs mRNA formation (*40, 41*). The structure further confirms the crucial role of U5 snRNA loop 1 in anchoring the 5'-exon and 3'-exon ends for catalysis of exon ligation. Thus, like the 5'SS GU dinucleotide and the branch point A nucleotide, recognition of the 3'SS AG dinucleotide is achieved through non-Watson-Crick RNA-RNA interactions stabilised by protein factors, explaining the conservation of these splice site sequences throughout eukaryotic evolution. An AU at the 5'SS and a AC at the 3'SS – a combination observed in the human minor spliceosome (*4*) and as a suppressor of 5'SS mutations in yeast (*30*) – could also be tolerated (**Fig. S6**), consistent with the major and minor spliceosome utilising a similar mechanism for 3'SS selection. Interestingly, although the structure of the RNA-based active site is remarkably conserved between the spliceosome and group II self-splicing introns (**Fig. S5**) (*18, 42, 43*), our P complex structure reveals that specific recognition of the 3'SS differs between the two splicing systems. Whereas in the group II intron the J2/3 junction interacts with the 3'SS (*44*), in the spliceosome the equivalent region in U6 snRNA (nucleotide A51) interacts instead with the 5'SS to stabilise the C* and P complex configuration of the active site (**Fig. 2B**) (*14*). Nonetheless, in both systems the 5'SS and the branch adenosine appear critical for both steps of splicing (**Fig. 2**) (*43, 45*), potentially explaining conservation of the 2'-5' linkage during evolution of both splicing systems.

Overall our P complex structure elucidates the mechanism of 3'-splice site selection, showing the crucial role of the excised lariat-intron in organising the active site and splice sites for exon ligation. The structure now completes our basic understanding of the two-step splicing reaction, showing how the dynamic protein scaffold of the spliceosome cradles and modulates a fundamentally RNA-based mechanism for splice site recognition during catalysis.

## Acknowledgements

M.E.W. prepared the sample, made EM grids, collected and processed EM data, carried out model building, and refined the structure. S.M.F. suggested the RNaseH method, prepared Prp22, and assisted data acquisition. W.P.G. was involved in the early stage of the project. M.E.W., S.M.F., and K.N. analysed the structure and drafted a manuscript. Mutagenesis and splicing assays were carried out by A.J.N. and C.M.N. The manuscript was finalized by M.E.W., S.M.F., and K.N. with input from all authors. K.N. initiated and coordinated the spliceosome project. We thank C. Savva, S. Chen, G. McMullan, J. Grimmett, and T. Darling for smooth running of the EM and computing facilities; the mass spectrometry facility for help with protein identification; and the members of the spliceosome group for help and advice throughout the project. We thank J. Löwe, V. Ramakrishnan, D. Barford, and R. Henderson for their continuing support and C. Plaschka, P. C. Lin, C. Charenton, C. J. Oubridge, and L. Strittmatter for critical reading of the manuscript. The project was supported by the Medical Research Council (MC_U105184330) and European Research Council Advanced Grant (SPLICE3D). M.E.W was supported by a Cambridge-Rutherford Memorial PhD Scholarship, S.M.F. was supported by EMBO and Marie Skłodowska-Curie fellowships. Cryo-EM maps will be available from the Electron Microscopy Data Bank and atomic models will be available from the Protein Data Bank.

## Materials and Methods

### Preparation and purification of P complex

Yju2-TAPS yeast was grown in a 120 L fermenter and splicing extract was prepared by the liquid nitrogen method essentially as previously described (*46, 47*) A DNA template for *in vitro* transcription was generated by addition of 3xMS2 stem loops to the 3'-end of a UBC4 pre-mRNA sequence containing a 20 nt 5'-exon, 95 nt intron, and 32 nt 3'-exon (*6, 48*). The RNA product was labelled at the 3' end with fluorescein-5-thiosemicarbazide (*49*). Prp22 dominant negative mutant (S635A) was expressed as a N-terminal calmodulin binding peptide fusion in yeast and purified as described previously (*34*). *In vitro* splicing reactions were assembled from pre-mRNA substrate pre-bound to MS2-MBP fusion protein as described (*50*), except that splicing extract for a 180 mL splicing reaction was supplemented with 1.8 mg of dominant-negative (S635A) mutant Prp22 protein (*51*). Reactions were incubated for 30 min at 23 ^o^C. The DEAH-box helicase Prp22 catalyses release of spliced mRNA from the spliceosome. Dominant-negative Prp22 therefore stalls complexes at the P complex stage. To remove spliceosomes without a docked 3'-exon, we then incubated the splicing reaction for 20 min with 5 μM of DNA oligonucleotide complementary to the 3'-exon (oligo sequence 5'-ATGAAGTAGGTGGAT-3') to induce cleavage of the 3'-MS2 tag by the endogenous RNaseH activity of the splicing extract. In contrast, P complexes with a docked 3'-exon were protected from RNaseH (*25*) and could be purified via MS2-MBP fusion protein (Fig. S1A). The reaction mixture was centrifuged through a 40% glycerol cushion in buffer A (20 mM HEPES, pH 7.9, 75 mM KCl, 0.25 mM EDTA). The cushion was collected and applied to amylose resin. After 15 h incubation at 4 ^o^C the resin was washed and eluted with buffer A containing 5% glycerol, 0.01% NP-40 and 12 mM maltose. Fractions containing spliceosomes were concentrated to 2 mg ml^−1^, dialysed against buffer A for 3 h, then used immediately for EM sample preparation. All proteins in the isolated complex were consistent with it being a second step spliceosome (Fig. S1). The complex contained mRNA and lariat intron with virtually no un-spliced pre-mRNA or lariat-intron intermediate, confirming its identity as complex P (Fig. S1). Smaller RNA species were not spliceosome-associated (Fig. S1) and likely represent the MS2 stem loops removed from spliceosomes that did not successfully undergo exon ligation.

### *In vivo* genetic screening of Prp8 mutants

Initial alanine screening of Prp8 was performed by the plasmid shuffling method. The Prp8 deletion-mutant strain SC261D8B1, carrying wild-type PRP8 on pRS316 (URA3, centromeric replication origin (*52*)) was transformed with mutant Prp8 on pRS314 (TRP1, centromeric replication origin (*52*)). Mutations were introduced using Kunkel mutagenesis, as described (*53*). Transformants were selected on plates lacking tryptophan and cells were then transferred onto plates containing 5-fluoro-orotic acid (5-FOA), to test cell growth after loss of the uracil plasmid. Viability was assessed visually after 3–5 days of incubation at 30 ^o^C. Transformants that survived on 5-FOA were colony purified and spotted onto YPD plates. Growth was then assessed after 3–5 days of incubation at 30 ^o^C. Mutations that confer lethality on 5-FOA or are viable but confer slow growth on YPD are indicated in Fig. S8.

### *In vitro* splicing

To assess splicing by spliceosomes containing Prp8 mutants that were lethal or conferred a slow growth phenotype, splicing extracts were prepared from merodiploid strains containing both an untagged, wild-type copy of Prp8 (on pRS316) and a Protein A-tagged, wild-type or mutant copy of Prp8 (on pRS314). Merodiploid yeast were grown in media lacking uracil and tryptophan to maintain both copies of Prp8 and extracts were prepared by the Dounce method, as described (*47*). For *in vitro* splicing a modified ACT1 pre-mRNA substrate was used (*53*) radiolabeled at the branch point. Following initial incubation under splicing conditions, reactions were diluted in IPP-150 buffer (10mM Tris-HCl pH 8; 150mM NaCl; 0.01% NP-40 substitute (Fluka)) and spliceosomes associated with the Protein A-tagged copy of Prp8 were immunoprecipitated with IgG-sepharose (GE). Following washing with IPP-150, the associated RNA species were phenol-extracted and analyzed on 6% denaturing polyacrylamide gels.

### Electron microscopy

For cryo-EM analysis the purified P complex sample was applied to R2/2 holey carbon grids (Quantifoil) coated with a ~7 nm homemade carbon film. Grids were glow discharged for 15 s before deposition of 3 μL sample, then incubated for 30 s and blotted for 2.5 s before vitrification by plunging into liquid ethane using an FEI Vitrobot MKIII operated at 100% humidity and 4 ^o^C. Grids were imaged in an FEI Titan Krios transmission electron microscope operated in EFTEM mode at 300 kV using the Gatan K2 Summit direct electron detector and a GIF Quantum energy filter (slit width 20 eV). Micrographs were collected automatically using EPU, collecting 2,384 movies with a defocus range of 0.2 – 3 μM at a nominal magnification of 105,000 (1.12 Å pixel^−1^). The camera was operated in counting mode with a total exposure time of 12 s fractionated into 20 frames, a dose rate of approximately 5 e^−^ pixel^−1^ s^−1^ and a total dose of 50 e^−^ Å^−2^ per movie.

### Image processing

Movies were corrected for movement using MotionCor2 (*54*), applying 5×5 patching and for initial processing applying dose-weighting to individual frames. CTF parameters were estimated using Gctf (*55*), before manual inspection of micrographs. We discarded 89 micrographs that had ice contamination or poor power spectra. From a subset of 216 micrographs, 16,551 particles were picked automatically using Gautomatch (Kai Zhang) and used to calculate reference-free 2D class averages in RELION 2.1 (*56, 57*). Three classes that represented the most common views of P complex were then used as templates for further picking with Gautomatch on all 2,295 good micrographs, yielding 206,529 particles. 3D refinement of all particles was performed with RELION using C* complex as a reference, yielding a reconstruction at an overall resolution of 3.9 Å. Particle polishing in RELION then improved the overall resolution to 3.8 Å.

To improve the map quality the polished particles were subject to 3D classification in RELION using 1.8° local angular sampling (Fig. S2). The 96,524 particles in the highest quality class were refined to 3.6 Å resolution, and then further classified with 1.8° local angular sampling with a soft mask that excluded U2 snRNP, U5 snRNP foot domain, Cwc22 NTD, U2/U6 helix II, U6 snRNA 5’stem, and the NTC proteins Prp19, Syf1, and Clf1 (C-terminus). This revealed the 3'SS docked and undocked states described in the main text. Refinement of the 48,617 particles in the 3'SS docked state produced a reconstruction at an overall resolution of 3.7 Å with a temperature factor of −82 Å^2^. Refinement of the 24,395 particles in the 3'SS undocked state produced a reconstruction at an overall resolution of 4.0 Å with a temperature factor of −80 Å^2^. Resolution is reported based on the gold-standard Fourier Shell Correlation as described (*58*).

### Structural Modelling

Model building was carried out in COOT (*59*). The majority of density could be accounted for by rigid-body fitting individual proteins from the yeast C* complex (*14*) and adjusting the fit where necessary. Because of the improved density quality in P complex for Slu7 and Prp17, new models were built for these regions: Slu7 was built entirely de novo into the density. Human Prp17 (*17*) was docked into the Prp17 density and mutated to the corresponding yeast sequences before manual adjustment. The CTD of Yju2, which was not modelled in C* complex, was built. The crystal structure of yeast Prp18 (*26*) was fitted into the density, and the conserved region not visible in the crystal structure was built. The α-finger of Prp8 (residues 1565-1610) was extended by 18 residues relative to C* complex. Residues 294-306 of the NTD of Prp22 were built into a pocket of the Prp8 reverse transcriptase domain. An updated model for the CTD of Ecm2 was docked into low-resolution density, based on a high-resolution map of C complex (unpublished data). The U5 snRNA model built for the C* complex was also replaced with U5 snRNA from B complex, which features the newly determined variable stem-loop II (*13*).

Coordinates for the core of P complex were refined in real space using phenix.real_space_refine in PHENIX (*60*) into the sharpened 3'SS-docked cryo-EM density, applying secondary structure, rotamer, base-pairing, base-stacking, and metal coordination restraints. For refinement of the lariat-intron 2'-5' linkage a custom set of restraints was adapted from the geometry of the 3'-5' phosphodiester RNA backbone. The peripheral proteins Prp22, Syf1, the C-terminal half of Clf1, U2 snRNP, the Cwc22 NTD, and the Prp19 module (comprising the Prp19 tetramer, Snt309 and the CTD of Cef1) were not refined. The final P complex model comprises 40 proteins, 3 snRNAs, and the intron and spliced exon (mRNA) products. Figures were generated with PyMOL (http://www.pymol.org) and UCSF Chimera (*61*).

### Data availability

The cryo-EM map will be deposited in the Electron Microscopy Data Bank. The coordinates of the atomic model will be deposited in the Protein Data Bank.

---

**Fig. S1.**
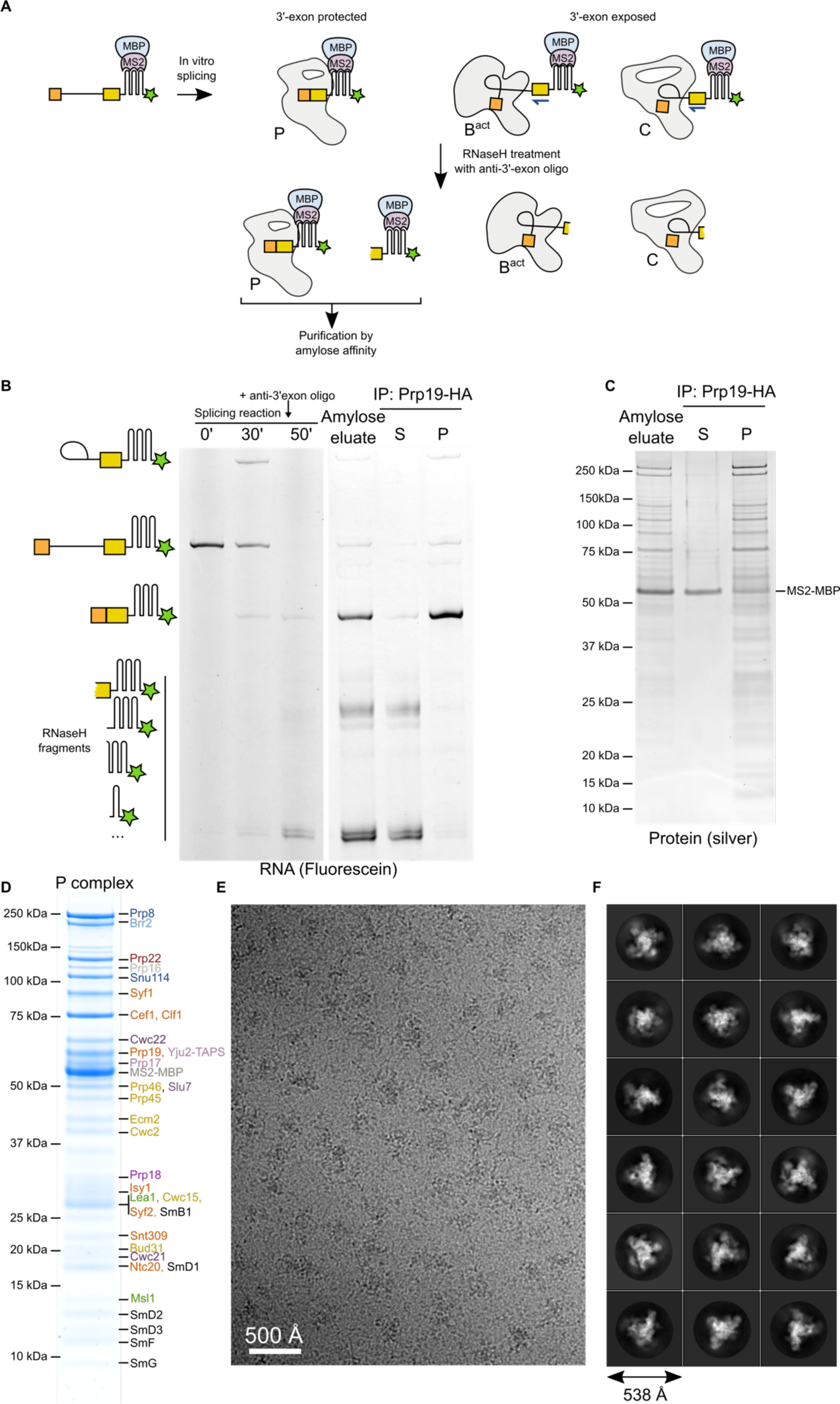
Purification and imaging of the P complex spliceosome. **A**, Schematic of the use of RNaseH protection to enrich for spliceosomes with a docked 3'-exon. Blue arrow represents anti-3'-exon DNA oligonucleotide, three hairpins represent 3xMS2 tag, green star represents 3' fluorescein label. **B**, The RNaseH strategy produces spliceosomes enriched for mRNA. Splicing was performed in Prp19-HA extract. Immunoprecipitation (IP) of the amylose eluate with anti-HA antibody shows that mRNA is precipitated (P) while smaller RNA species are in the supernatant (S), showing these are not associated with activated spliceosomes. **C**, SDS-PAGE showing total proteins (stained with silver) associated with the amylose eluate and with the P and S fractions from Prp19-HA IP. The smaller RNA species in S are associated with MS2-MBP while all spliceosomal proteins are in the precipitate. **D**, SDS-PAGE showing total proteins (stained with Coomassie Instant Blue) in the P complex sample used for cryo-EM analysis (concentrated amylose eluate). Labelled bands were excised and identified by mass spectrometry and are coloured by subcomplex identity (blue, U5 snRNP; green, U2 snRNP; orange, NTC; yellow, NTR; purple, splicing factors; black, Sm proteins; grey, not modelled). **E**, Representative cryo-EM micrograph of vitrified P complex. **F**, Representative 2D class averages calculated in RELION for P complex.

**Fig. S2.**
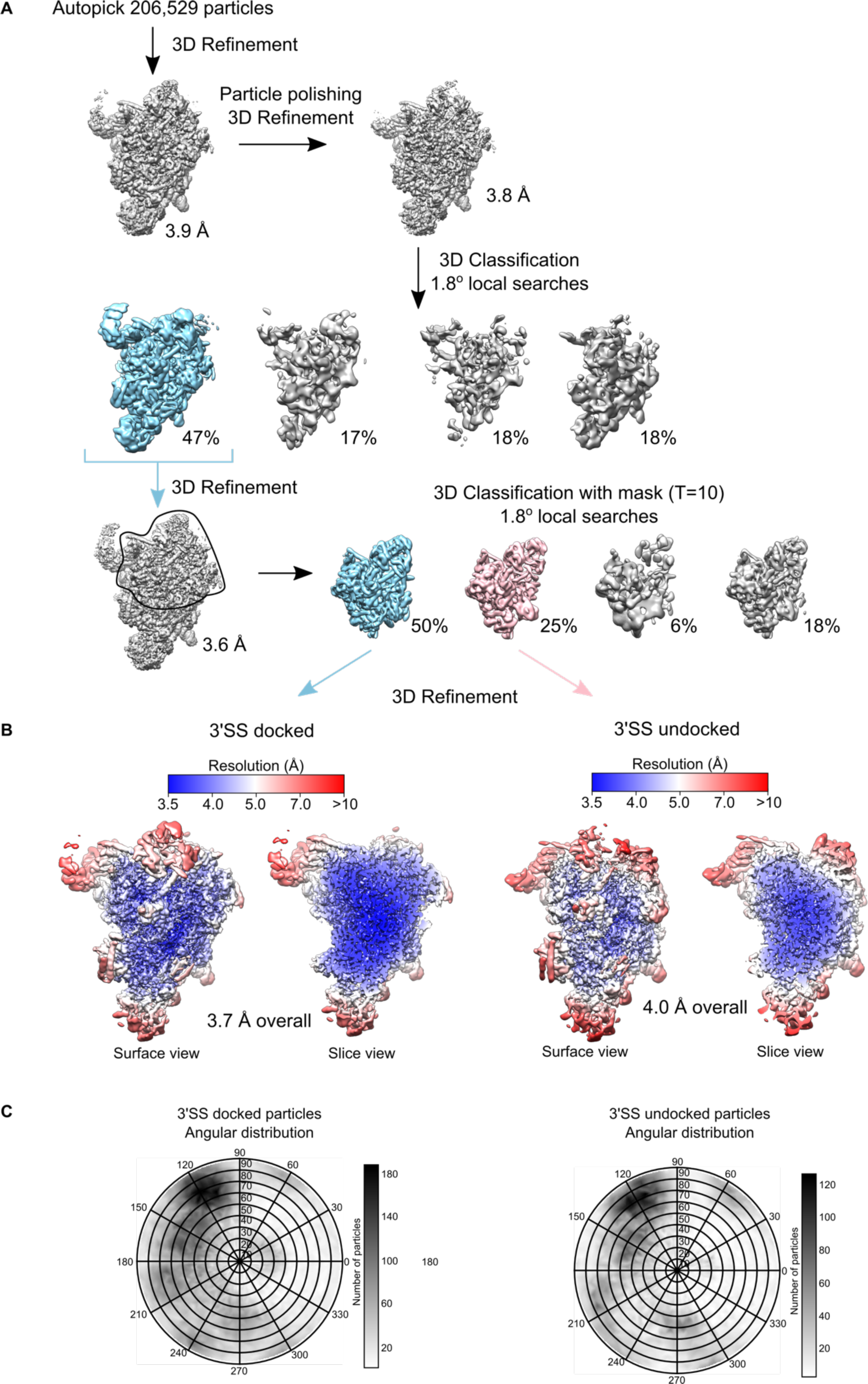
Cryo-EM data processing. **A**, Classification scheme. All autopicked particles were refined and polished before 3D classification with local angular searches. The most populated, high-resolution class was refined before focused classification with a mask around the upper half of the core structure (indicated), producing classes corresponding to the 3'SS docked (blue) and undocked (pink) state of P complex. Note that global (unmasked) classification of the entire data set also produced the docked/undocked states in a ~1:1 ratio but at a lower overall resolution when refined. **B**, Surface and central slices through the cryoEM reconstructions of the 3'SS docked/undocked states, coloured by local resolution as calculated in RELION. **C**, Orientation distribution plot for particles contributing to the 3'SS docked/undocked 3D reconstructions.

**Fig. S3.**
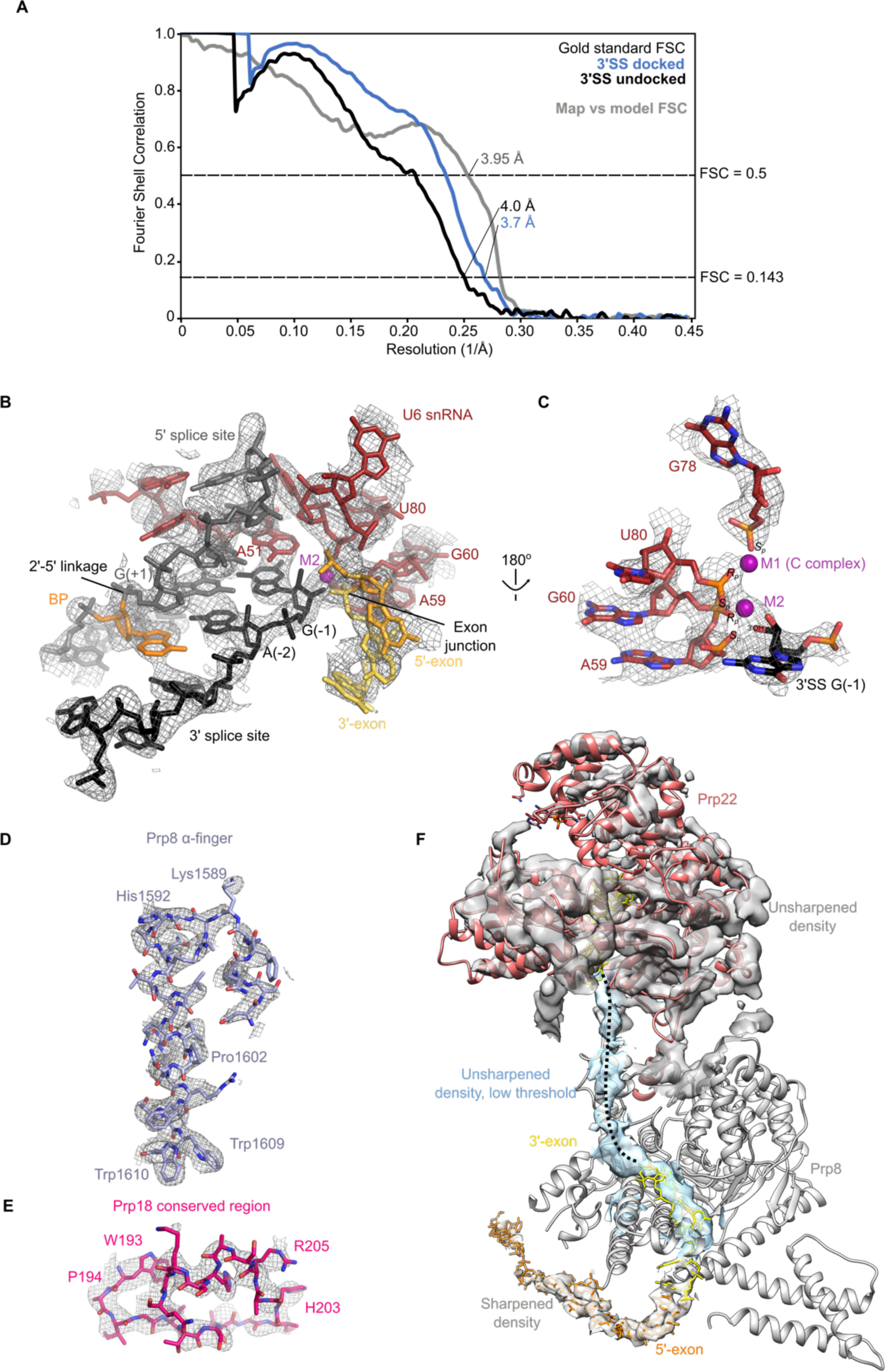
Examples of Cryo-EM density and built models. **A**, Gold-standard Fourier shell correlation (FSC) for the 3'SS docked/undocked 3D reconstructions, as determined by post-processing with a soft mask in RELION; Map-vs-model FSC for the refined atomic coordinates against the 3'SS docked sharpened cryo-EM density. **B-F** show example density from the 3'SS docked map. **B**, Substrate RNA bound at the catalytic RNA centre of P complex. **C**, Coordination of catalytic metals. M2 was modelled into density; no density is visible for M1 and the position from C complex is indicated. **D**, Prp8 α-finger region. **E**, The conserved region of Prp18. **F**, Cryo-EM density for the 3'-exon leading to Prp22. The same map is shown sharpened for the 5'-exon and start of 3'-exon, unsharpened for Prp22 (due to the lower local resolution in this region) and unsharpened at low threshold (blue) for the weak density interpreted as the 3'-exon connecting to Prp22.

**Fig. S4.**
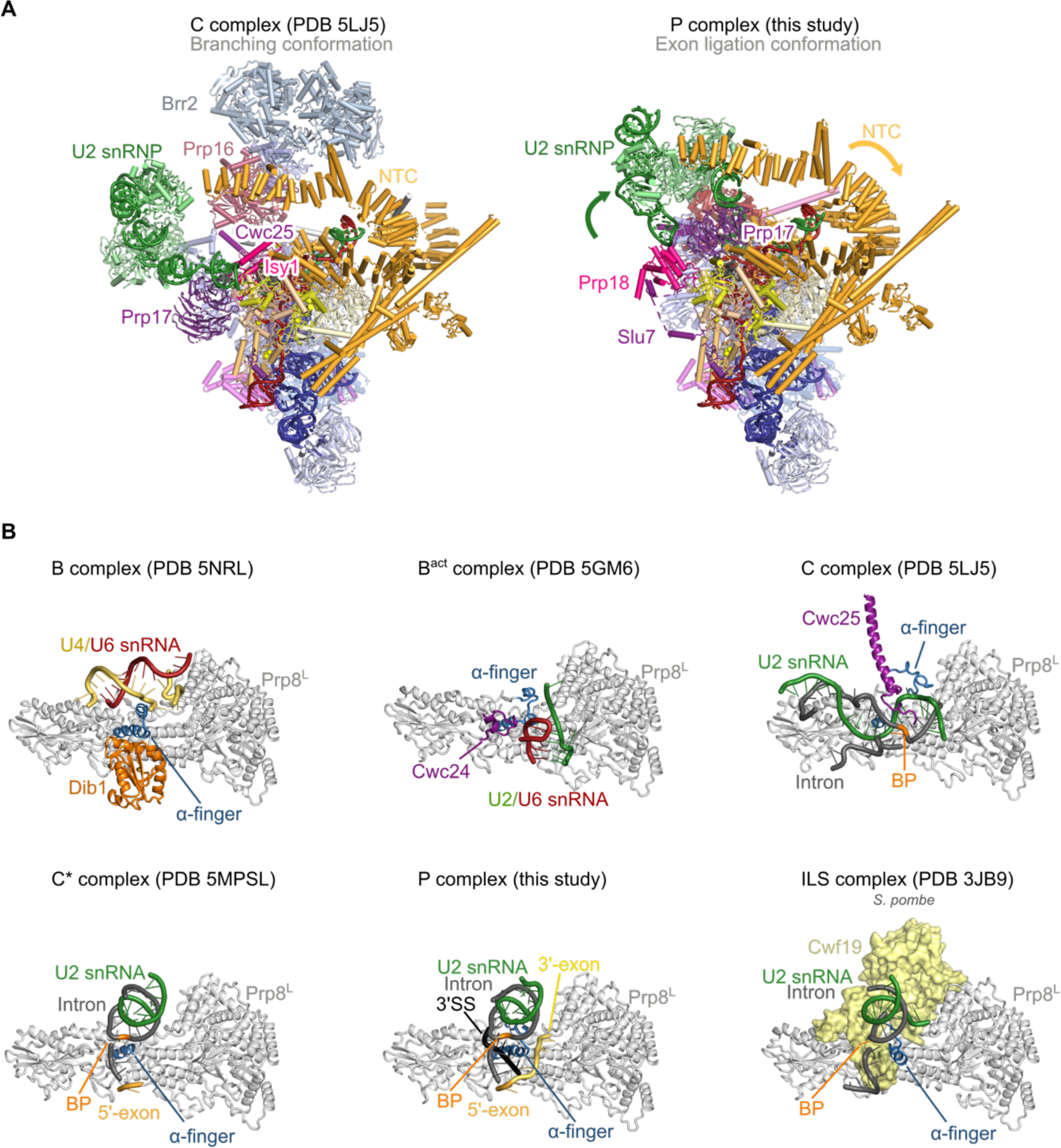
Spliceosome remodeling. **A**, Conformational changes between branching (complex C; PDB ID 5LJ5) and exon ligation (complex P; this study) conformations of the spliceosome. **B**, Remodeling of the Prp8 α-finger through the splicing cycle. The Prp8 α-finger (residues 1577-1610; 1527-1562 for *S. pombe*) is shown in blue for various yeast spliceosomal complexes. The α-finger makes context-dependent contacts with various components of the splicing machinery.

**Fig. S5.**
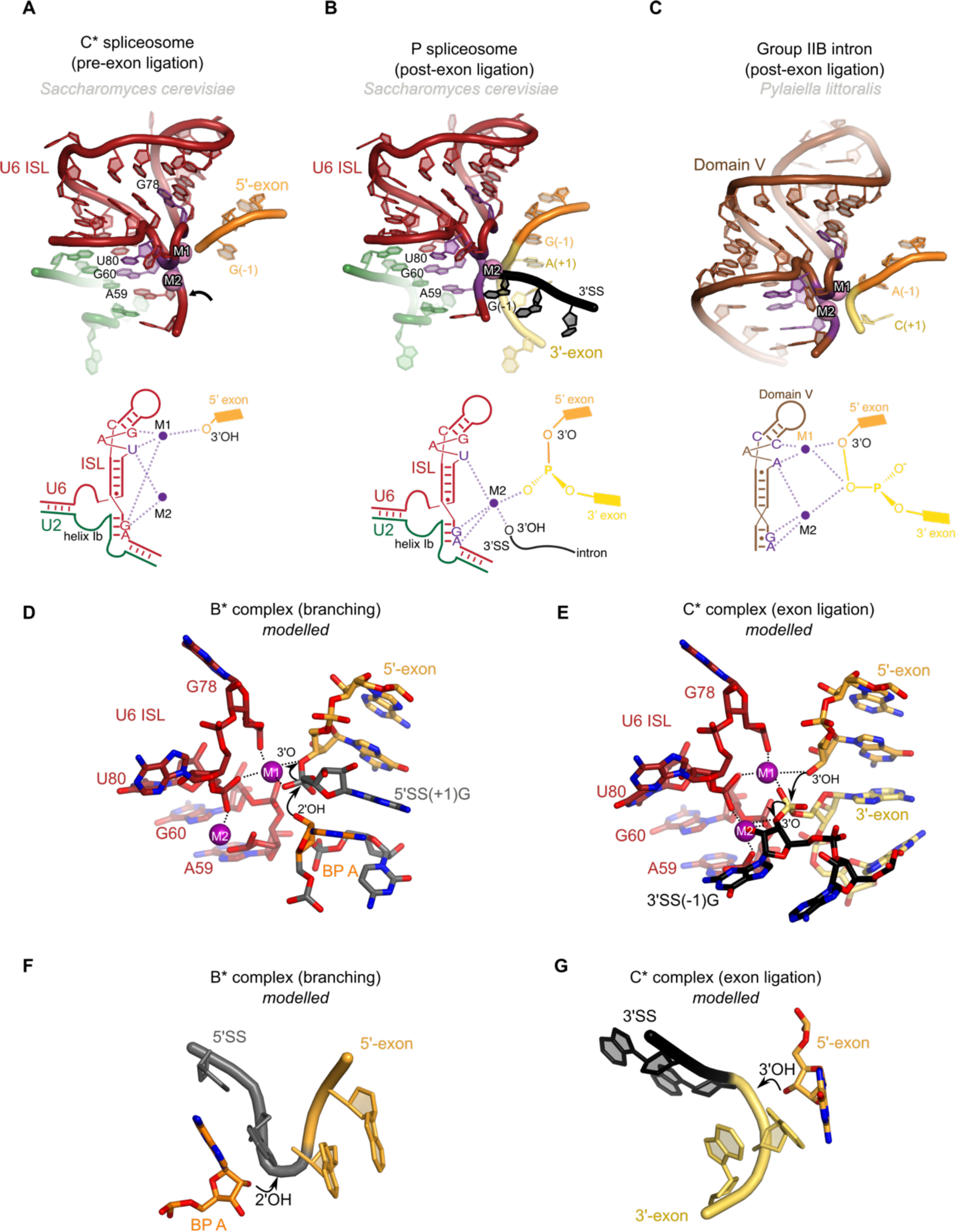
Metals in the RNA core of the P complex. **A**, Structure and schematic representation of the active site of the C* complex trapped before exon ligation (PDB 5MPS). Note the proximity of the cleaved G(-1) 3'-hydroxyl to M1. Density for M2 was weakly visible in C* and was coordinated by phosphates from G60 and U80. **B**, Structure and schematic representation of the active site of P complex. Note that M2 is now coordinated by phosphates from A59, G80, and U80 and is also interacting with both the newly formed mRNA junction and the cleaved 3'OH from the 3'SS. Note that to accommodate the docked 3'-exon, the phosphate backbone between C58 and A59 moves to position the A59 phosphate for interacting with M2. **C**, Structure and schematic of the RNA core of a group IIB intron in a post-catalytic configuration, showing the ligated exons (PDB 4R0D). Residues that position the catalytic metals are shown in magenta. **D, E**, The inferred structure of the spliceosome active site during branching and exon ligation from the same view. Structures of B* and C* are modelled based on C complex (PDB 5LJ5) and P complex (this study) by fixing ribose and base atoms and regularizing the bonding geometry over the scissile phosphate for the pre-reaction state. Note that the scissile phosphate is in the same position relative to M1 for branching and exon ligation. For B* complex M1 and M2 are modelled based on PDB 5GMK. For C* complex M2 is positioned as in P complex; M1 was not visible in P so its position is based on C complex and is for illustrative purposes. **F, G**, The structure of the 5'SS and 3'SS before branching and exon ligation respectively. Reactants are shown from the same view relative to U6 snRNA. In B* the 5'SS has a tight kink to expose the scissile phosphate; in C* the 3'SS has a less pronounced kink. Note that the nucleophiles for each reaction attack the SS from opposite sides.

**Fig. S6.**
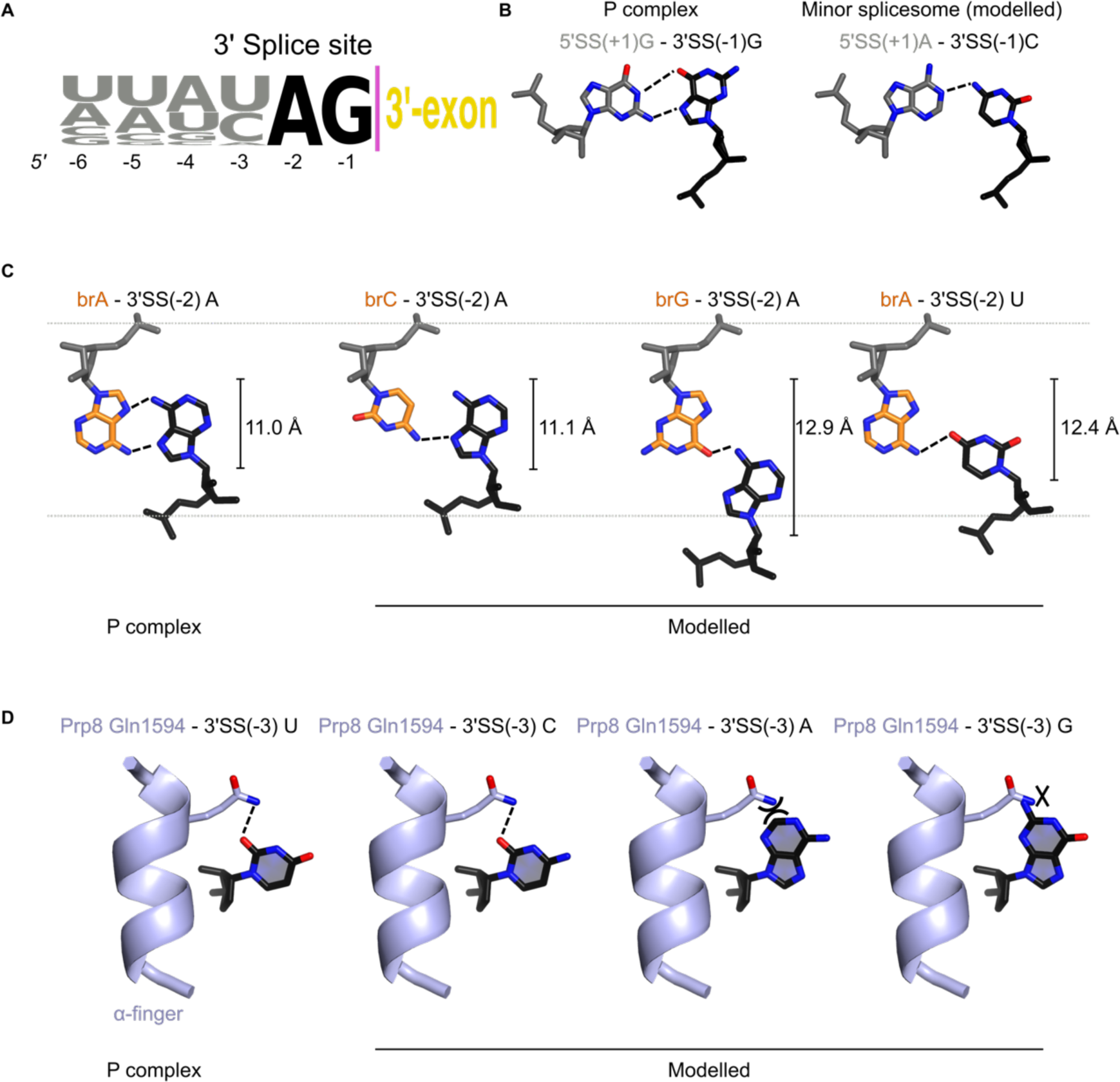
Non-Watson-Crick interactions mediate 3' splice site recognition. **A**, The consensus 3' splice site sequence in *S. cerevisiae*. The 3' end of all *S. cerevisiae* introns were aligned and used to create a sequence logo (weblogo.berkeley.edu). The invariant AG dinucleotides are highlighted, and the splice junction is indicated with a purple line. **B**, The cis Watson-Crick/Hoogsteen base pair that mediates recognition of 3'SS(-1)G by the 5'SS(+1)G (left). A similar interaction may operate in the minor spliceosome, where 25% of introns start with A and end with C (right). Note that the A-C base pair has not been observed experimentally, so is here modelled by mutation in COOT. **C**, The trans Hoogsteen/Hoogsteen base pair that mediates recognition of the 3'SS(-2)A by the branch point adenosine (brA) (left). The branch is highlighted in orange. Right, the effects of various branch and 3'SS mutants on this pairing, modelled based on observed examples highlighted in (*62*). The C1'-C1' distances are indicated, also taken from (*62*). Branch C has a minor step 2 defect, while branch G and 3’SS(-2)U have stronger defects, correlating with increased distortion relative to P complex. **D**, Recognition of the 3'SS(-3) pyrimidine by glutamine 1594 (Gln1594) of the Prp8 α-finger. In our P complex structure 3'SS(-3) is U (left). Cytidine can be tolerated in the same position, whereas a purine would have a minor clash with the alpha finger (right).

**Fig. S7.**
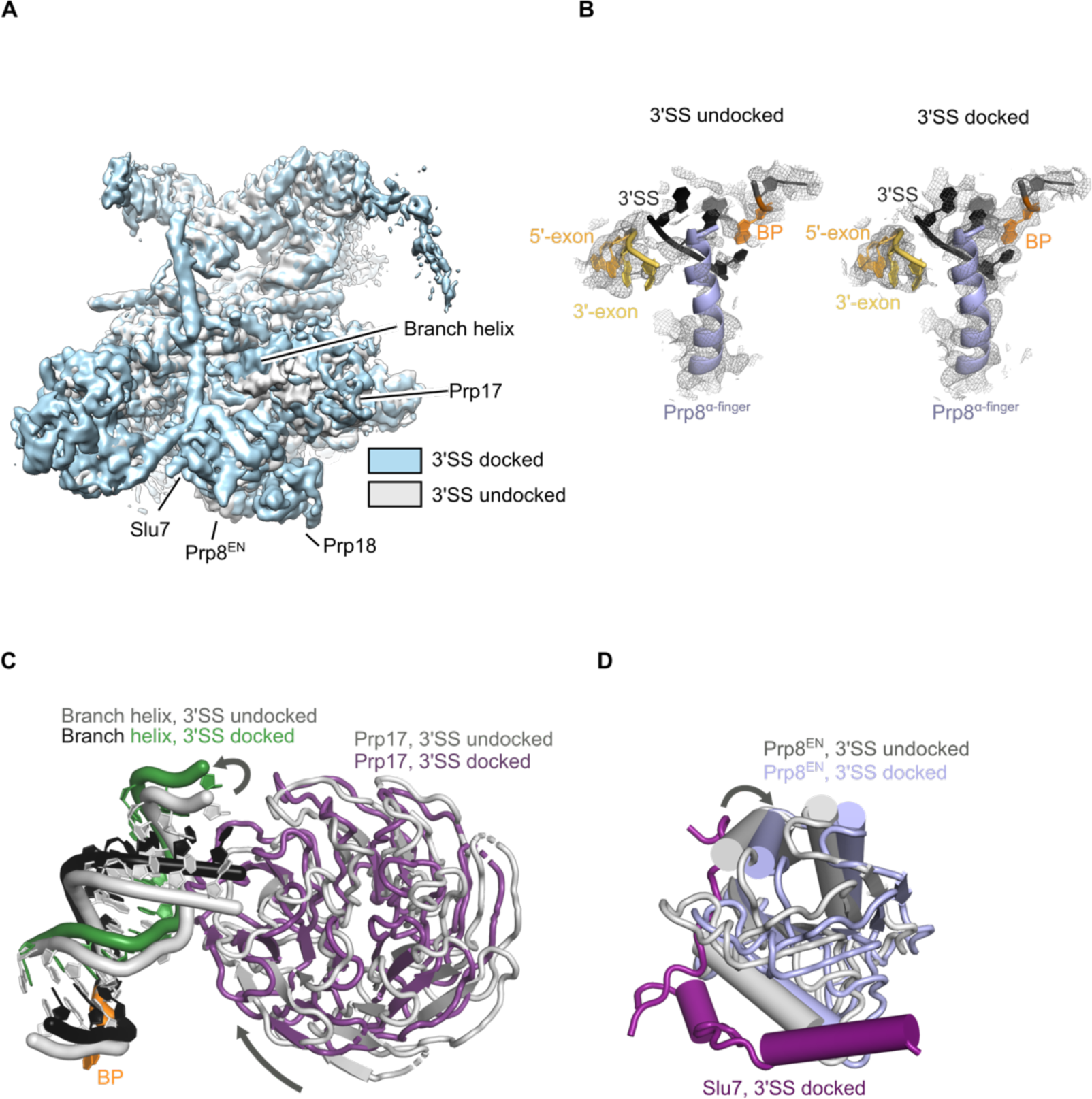
The docked and undocked states of P complex. **A**, Cryo-EM density low-pass filtered to 5 Å for the 3'SS undocked (grey) and docked (blue) states was superimposed. Regions that move or change occupancy are labelled. **B**, Sharpened density around the active site for the undocked and docked states. BP, branch point adenosine; 3'SS, 3' splice site. **C,D** Atomic models from the docked state (coloured) were fitted into the undocked state cryo-EM density. **C**, the branch helix and Prp17 pivot around the branch point. **D**, Prp8 endonuclease domain (EN) rotates between the two conformations, with the docked conformation associated with binding of Slu7.

**Fig. S8.**
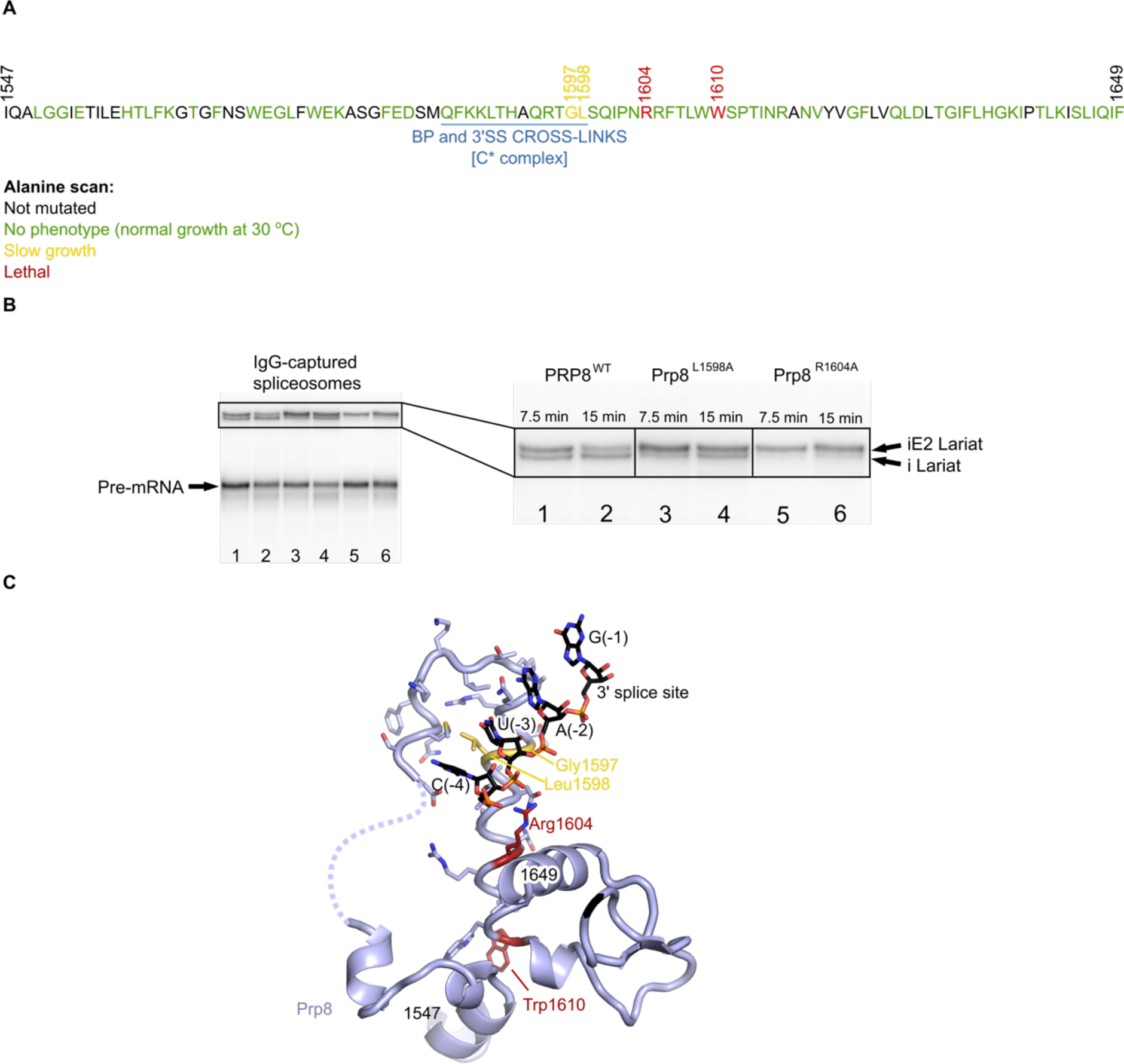
Characterisation of Prp8 α-finger mutations. **A**, Alanine-scanning mutagenesis of Prp8 residues 1547-1649. All residues except those in black were mutated to alanine. Those that produced no phenotype are shown in green, residues that produced slow growth in yellow, and lethal mutations in red. **B**, In vitro splicing with extract prepared from merodiploid *S. cerevisiae* expressing both WT Prp8 and either WT, L1598A, or R1604A mutant Prp8 with a protein A tag. Protein A-immunoprecipitated spliceosomes were analysed for RNA content by denaturing PAGE. L1598A, which causes slow growth, and R1604, which is lethal, both have defects in exon ligation. **C**, Structure of the Prp8 α-finger from P complex with slow growth or mutant alleles highlighted by the same colouring scheme. R1604 contacts the backbone of the 3'SS, while W1610 has a structural role in a hydrophobic pocket. No spliceosomes could be immunoprecipitated from extract with W1610A. Gly1597 contacts 3'SS(-3); any mutation in this position would cause a steric clash. Leu1598 forms a hydrophobic interface with the N-terminal helix of the α-finger that probably aids formation of the correct structure. i Lariat, intron-lariat; iE2 Lariat; lariat-intron-3'-exon intermediate.

**Table S1.**
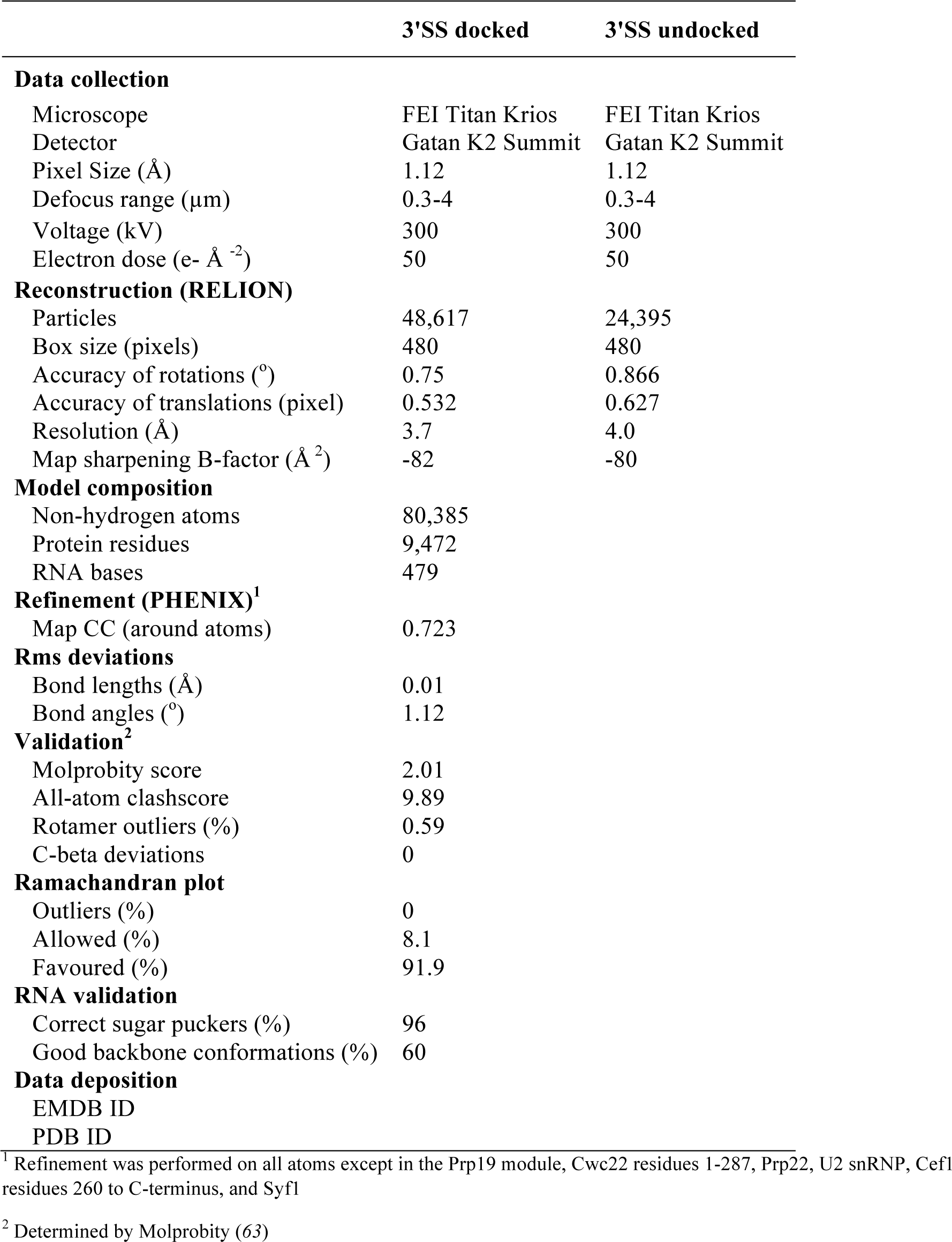
Cryo-EM data collection and refinement statistics

|  | 3'SS docked | 3'SS undocked |
| --- | --- | --- |
| **Data collection** |  |  |
| Microscope | FEI Titan Krios | FEI Titan Krios |
| Detector | Gatan K2 Summit | Gatan K2 Summit |
| Pixel Size (Å) | 1.12 | 1.12 |
| Defocus range (µm) | 0.3-4 | 0.3-4 |
| Voltage (kV) | 300 | 300 |
| Electron dose (e<sup>-</sup> Å<sup>-2</sup>) | 50 | 50 |
| **Reconstruction (RELION)** |  |  |
| Particles | 48,617 | 24,395 |
| Box size (pixels) | 480 | 480 |
| Accuracy of rotations (°) | 0.75 | 0.866 |
| Accuracy of translations (pixel) | 0.532 | 0.627 |
| Resolution (Å) | 3.7 | 4.0 |
| Map sharpening B-factor (Å<sup>2</sup>) | -82 | -80 |
| **Model composition** |  |  |
| Non-hydrogen atoms | 80,385 |  |
| Protein residues | 9,472 |  |
| RNA bases | 479 |  |
| **Refinement (PHENIX)<sup>1</sup>** |  |  |
| Map CC (around atoms) | 0.723 |  |
| **Rms deviations** |  |  |
| Bond lengths (Å) | 0.01 |  |
| Bond angles (°) | 1.12 |  |
| **Validation<sup>2</sup>** |  |  |
| Molprobit score | 2.01 |  |
| All-atom clashscore | 9.89 |  |
| Rotamer outliers (%) | 0.59 |  |
| C-beta deviations | 0 |  |
| **Ramachandran plot** |  |  |
| Outliers (%) | 0 |  |
| Allowed (%) | 8.1 |  |
| Favoured (%) | 91.9 |  |
| **RNA validation** |  |  |
| Correct sugar puckers (%) | 96 |  |
| Good backbone conformations (%) | 60 |  |
| **Data deposition** |  |  |
| EMDB ID |  |  |
| PDB ID |  |  |

<sup>1</sup> Refinement was performed on all atoms except in the Prp19 module, Cwc22 residues 1-287, Prp22, U2 snRNP, Cef1 residues 260 to C-terminus, and Syf1
<sup>2</sup> Determined by Molprobit (63)

**Table S2.**
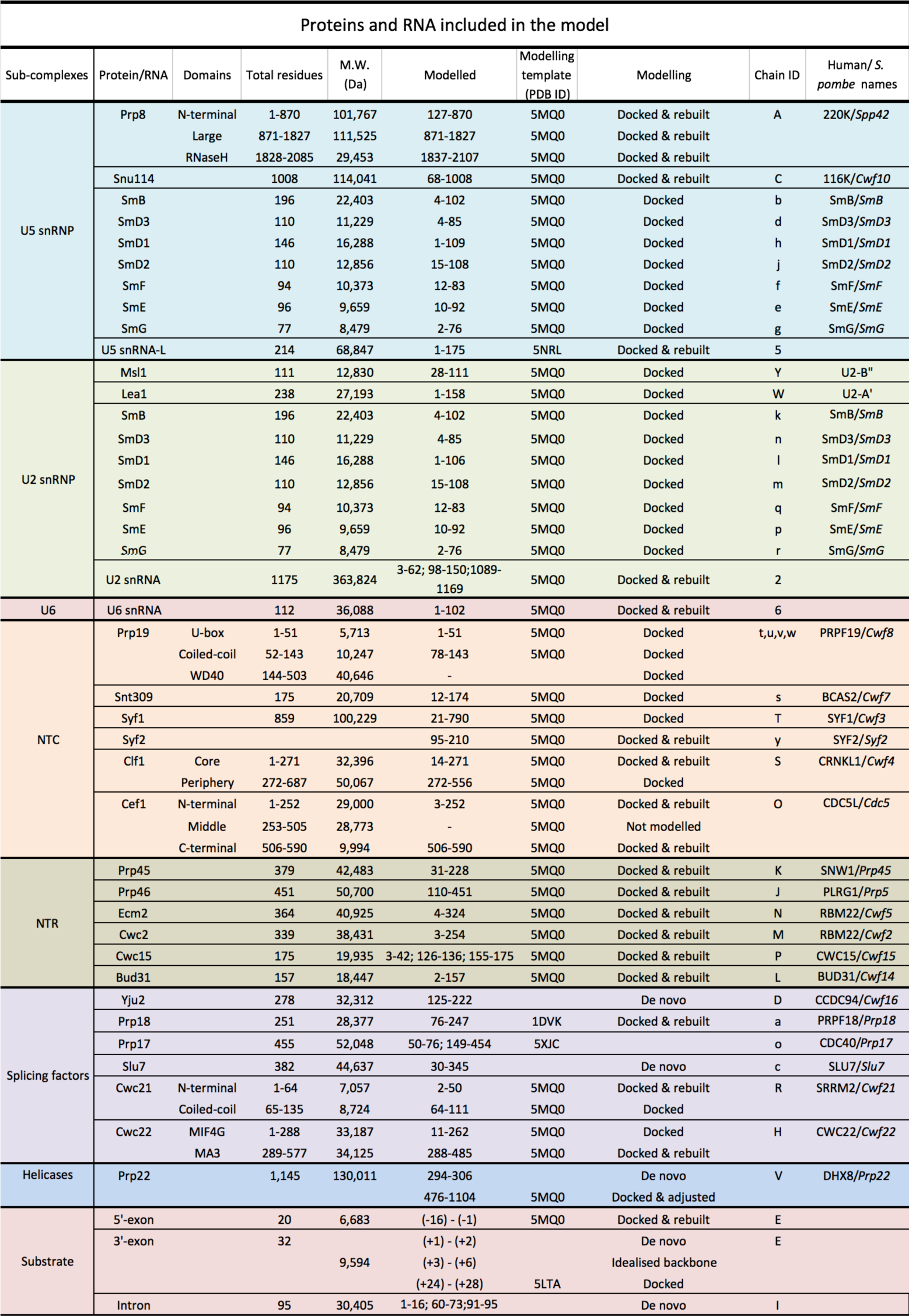
Summary of components modelled into the P complex map

